# Multiday rhythms modulate human heart rate: an observational study in healthy adults

**DOI:** 10.64898/2026.03.01.708870

**Authors:** Rochelle I. De Silva, Rachel E. Stirling, Jodie Naim-Feil, Shivam Puri, Elizabeth Paratz, Philippa J. Karoly

**Affiliations:** Department of Biomedical Engineering, University of Melbourne, Melbourne, Australia; Graeme Clark Institute, University of Melbourne, Melbourne, Australia; Department of Medicine, St. Vincent’s Hospital Melbourne, University of Melbourne, Melbourne, Australia

**Author notes:** Corresponding author Philippa J. Karoly, Full address: Building 261/203 Bouverie Street, Carlton, VIC 3053, Australia. Authors contributed equally to this work (Rochelle I. De Silva, Rachel E. Stirling).

**Keywords:** infradian, biological rhythms, cardiac, autonomic, wearables

## Abstract

**Background:** Chronobiology research has historically focused on circadian rhythms; however, longer infradian rhythms are prevalent in human physiology and may have important implications for health and wellbeing. Previous studies have identified widespread infradian rhythms across human physiology, often in the context of hormonal regulation and disease. Despite growing evidence of their ubiquity, the mechanisms, significance, and clinical relevance of these rhythms remain poorly understood, largely due to lack of longitudinal datasets and robust detection methods. The emergence of new wearable technologies enables rich, continuous data capture within individuals, allowing physiological rhythms to be studied at scale.

**Methods:** This study analyzed a cohort of healthy, young adults (N=623), with up to four years of wearable and questionnaire data collected through the University of Notre Dame’s (USA) NetHealth project. Participants who recorded at least three months of continuous (>80% adherence) heart rate data were included and significant infradian rhythms were identified using wavelet analysis. Unsupervised non-negative matrix factorization was performed to cluster similar wavelet power spectrum distributions. Individuals’ heart rate rhythms were compared to known environmental cycles (day-of-week, lunar, seasonal) and considering demographics and social networks. A second, smaller cohort (N=70) with heart rate and menstrual timing were included to analyze the interplay of hormonal regulation on monthly cycles. Multinomial logistic regression, and statistical tests (i.e., one-way ANOVA) were applied to quantify the effects of environmental, behavioral and demographic factors on heart rate rhythms.

**Findings:** Significant infradian rhythms of heart rate were detected in 69.7% (365/523) of the cohort and 35.9% (188/523) had two or more rhythms. Annual, biannual and 10-week rhythms were the most common. Within the 4–45-day band, individuals clustered into four multiday chronotypes based on dominant periodicities in their wavelet power spectra: weekly (∼7 days), shorter-monthly (∼25 days), longer-monthly (∼35 days), and multi-month (>35 days). Heart rate rhythms were influenced by environmental cycles (day-of-week and seasonality) but were not tightly correlated to external cues. Additionally, heart rate rhythms were synchronized to the menstrual cycle in most menstruating females, although monthly rhythms were also observed in males and menopausal women.

**Interpretation:** The prevalence of infradian, or multiday heart rate rhythms in healthy young people motivates further scientific investigation to understand the mechanisms of these rhythms and their potential association with autonomic function, and risk of disease or disease-specific symptoms. Characterizing physiological rhythms can drive new insights into how multiscale fluctuations modulate disease symptoms across neurological, psychiatric, and broader health conditions.

## Introduction

Circadian and longer, infradian rhythms are pervasive in the natural world, shaping reproduction, development, activity and behavior across marine and terrestrial organisms. Circadian biological rhythms are well-documented in humans and other organisms and are entrained by the external environment via light-dark cues from the solar day. These rhythms can persist in the absence of environmental cues (i.e., free-running) driven by endogenous, self-sustained oscillators. Unlike circadian rhythms, longer biological rhythms remain less widely characterized; however, infradian rhythms in humans have been observed across timescales, including circaseptan (7-day, about-weekly)^1–3^, circatrigintan (30-day, about-monthly)^4–6^, and seasonal or circannual (1-year, about-yearly)^7–9^. Examples of these infradian rhythms include mammalian growth cycles of enamel and bone deposition^10,11^, human heart rate^12,13^, and blood pressure^2,3,14^. Moreover, studies conducted in isolation report longer rhythms in heart rate, blood pressure, and hormone secretion can persist in the absence of zeitgebers (i.e., external environmental cues)^1,2^. One study documented circaseptan rhythms in heart rate and blood pressure in a woman living in social isolation over a 14-week period^2^. More recent studies in cohorts with chronic brain implants found longer ‘multiday’ rhythms of epileptic activity independent from environmental cues^15–17^. Although individual studies remain limited in size or scope, collectively there is an emerging body of evidence across immunology^18^, cardiology^13,19^, and neurology^12^ showing longer rhythms in humans that appear to follow free-running cycles with periods from weeks to months.

Several mechanisms have been proposed as potential drivers of infradian rhythms observed in humans. While fixed calendar-based (e.g. weekday^1^) or environmental (e.g. lunar) cycles^6^ may explain some observations, longer rhythms are also hypothesized to arise from internal mechanisms, including, hormonal fluctuations^8,9,20^, immune system fluctuations^18,21^, metabolic processes^22,23^, or autonomic regulation^12^. The menstrual cycle often dominates perceptions of monthly rhythms; yet, men also exhibit cyclical patterns on a monthly scale^4,12^. This is particularly evident in neurological cohorts where similar monthly fluctuations in brain and seizure activity are observed for men, women and children^15,16,24,25^. Furthermore, neurological rhythms have individual-specific periods ranging from weekly to seasonal^24^; therefore, mechanistic theories targeted at one cycle length cannot account for the striking prevalence nor heterogeneity of rhythms observed across central and autonomic nervous activity. Indeed, 60% of adults with epilepsy show infradian rhythms of brain activity, with similar proportions having weekly, fortnightly, monthly or longer cycles^26^. Outside the brain, a study of over 300 patients with cardiovascular disease found many had annual (78%), 7-day (26%) and 3-monthly (22%) rhythms in heart rate, although monthly cycles were rare^13^. A smaller cohort study using wearables found weekly and monthly heart rate rhythms were common in healthy adults^12^. Despite increasing evidence that infradian rhythms are a feature of human brain and heart activity, these rhythms have never been systematically identified in the wider population and consequently their underlying mechanisms, functional significance, and clinical relevance remain largely unknown.

Recent technological advances to enable continuous physiological monitoring and large-scale population data provide a unique opportunity to address whether infradian rhythms are endogenous or environmental. Wearable signals to track peripheral oscillatory systems are increasingly validated as proxies for less accessible central or hormonal oscillatory dynamics^27,28^, and offer longitudinal insights into coupled physiological rhythms between the heart and brain^12,29^ This study aimed to investigate long-term rhythms of heart rate in healthy adults, characterizing their prevalence, heterogeneity, and environmental influences, using a large database of wearable recordings^30^. The majority had heart rate rhythms with cycle periods ranging from weekly, monthly to longer multi-month or seasonal periods. Establishing normative patterns on infradian rhythms in healthy adults is a critical step toward identifying their governing drivers and understanding how multiday fluctuations can arise in disease symptoms across neurological^15,24^, psychiatric^20,31^, and broader health conditions. Advancing this understanding could reshape current models of physiology and drive novel applications in disease prediction, monitoring, and treatment.

## Results

This study analyzed data from the University of Notre Dame’s NetHealth project^32^ (Indiana, U.S.A.) in 623 college students who recorded continuous physiology via a smartwatch (daily heart rate, sleep, physical activity) alongside other surveys (including basic demographics, circadian preference, social networks) between August 2015 and May 2019. The full dataset and accompanying documentation are accessible at https://sites.nd.edu/nethealth/. The primary aim of the current study was to assess the prevalence of infradian rhythms in heart rate, with a focus on weekly to monthly cycles i.e., between 4 – 45 days (henceforth referred to as ‘multiday rhythms’).

Heart rate data was recorded from 623 participants, from which 100 were excluded due to insufficient data (<3 months) for cycle detection, leaving 523 (263 female). The mean recording duration across these individuals was 634.4 days (minimum: 93 days; maximum: 1364 days; standard deviation, SD=439.2 days). Further data characteristics are provided in Table 1 and Supplementary Table 1. Figure 1 shows recordings from two participants, with significant rhythms in their daily heart rate, detected using spectral analysis. Across the cohort, the detected multiday rhythms were compared across known environmental factors, including day-of-week, lunar and seasonal cycles and – for a different subset of women – the menstrual cycle.

**Figure 1.**
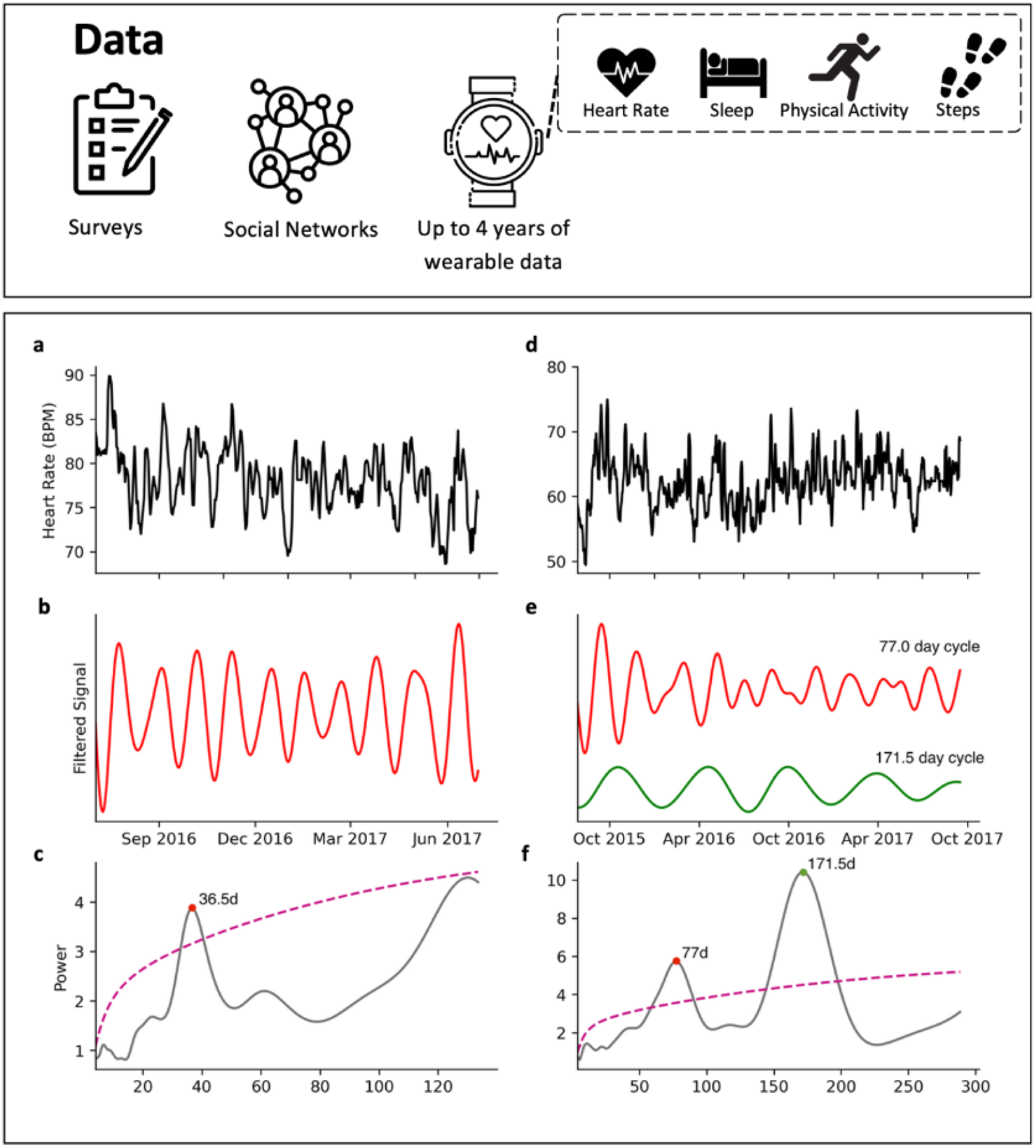
Schematic of the preprocessing and multiday cycle detection steps. Survey, social network and wearable smartwatch data (up to 4 years) were provided in the NetHealth project dataset. Heart rate data and multiday cycles are shown for two different individuals: example 1 (**a-c**) and example 2 (**d-f**). **a,d**: Daily average heart rate (y-axis) in beats per minute, smoothed with a 4-day moving average filter, shown over 1 year in example 1 and over 2 years in example 2. **b,e**: A graphical representation of the significant multiday cycles identified in the wavelet periodogram. The original daily average heart rate signal was bandpass filtered to produce the representation, with the passband including frequencies within the full-width-half-maximum of the corresponding peak. **c,f**: Wavelet periodogram for each example at different scales (x-axis). Significant cycle periods (distinct peaks) are labelled with colored dots.

**Table 1.** Data characteristics.

| Number of Participants |  |  |
| --- | --- | --- |
| Sex |  |  |
| Male |  | 317 |
| Female |  | 306 |
| MEQ Chronotypes |  |  |
|  | Summer 2015 | Winter 2016 |
| Definite Evening | 5 | 11 |
| Moderate Evening | 71 | 132 |
| Intermediate | 342 | 379 |
| Moderate Morning | 57 | 24 |
| Definite Morning | 1 | 1 |
| (No Response) | 147 | 76 |

### Multiday rhythmicity in human heart rate

Overall, 69.7% (365/523) had at least one significant rhythm of ≥4 days and 35.9% (188/523) of people had two or more, with significant rhythms determined from peaks in individuals’ wavelet periodograms (see Figure 1 and Supplementary Figure 1). This study first considered multiday heart rate rhythms from 4 – 45 days, which have been previously documented in neurological cohorts^17,26^. Similarities in wavelet periodogram distributions across the cohort of individuals with at least 135 days of heart rate data and significant rhythms (n=353) were identified using an unsupervised component decomposition (non-negative matrix factorization; NMF). Four common distributions, referred to as *multiday chronotypes*, were learned from the data (Figure 2), with four components chosen due to highest feature cross-correlation values (Supplementary Figure 2). These components had peaks centered around 7-days (weekly, n=70), 25-days (shorter monthly, n=97), 35-days (longer monthly, n=157), and longer (multi-month, n=29) periods.

**Figure 2.**
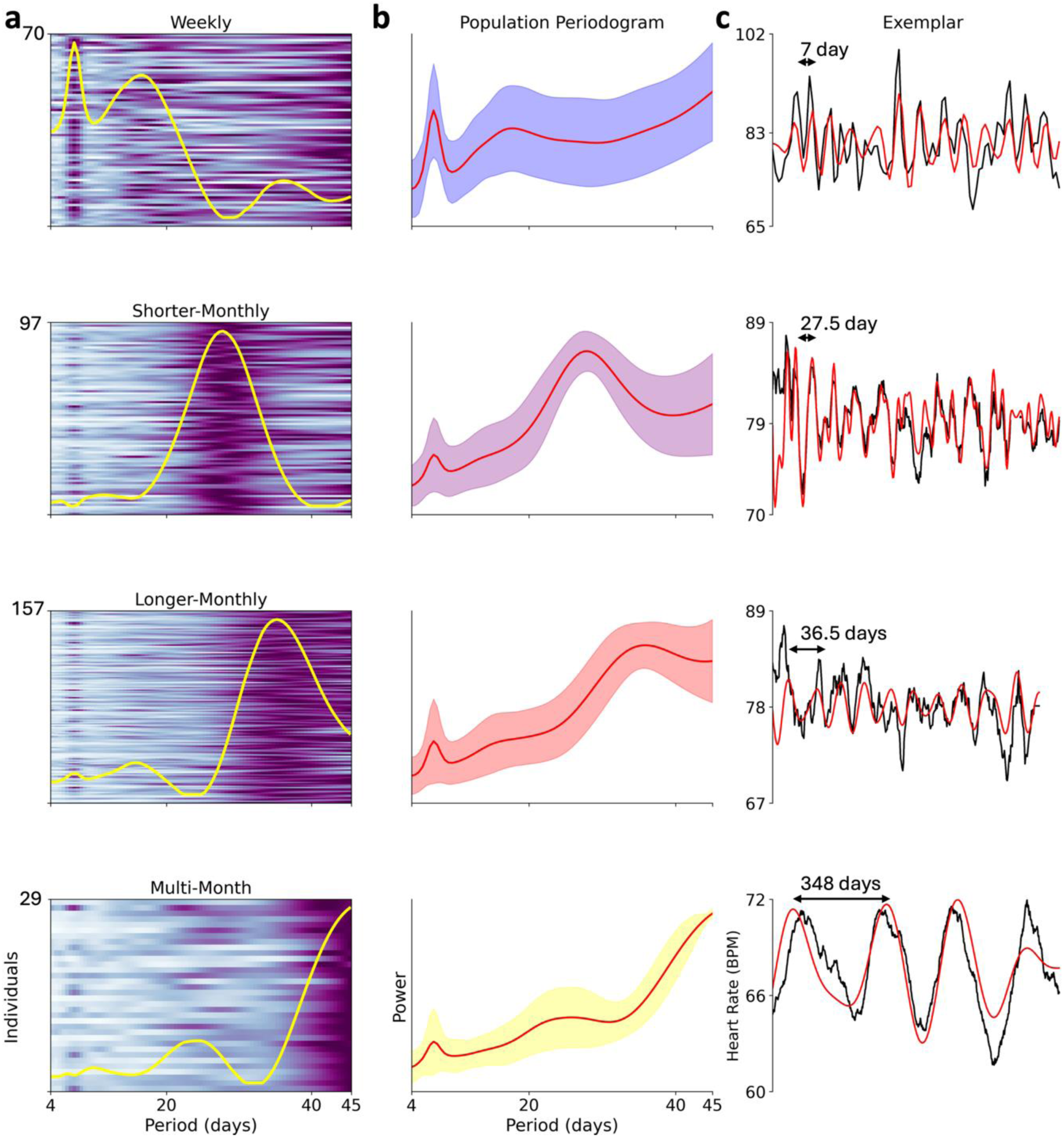
Multiday chronotypes identified from heart rate cycles. **a)** Heatmaps visually represent patterns of cyclic heart rate fluctuations within different chronotypes. Each row represents 1 individual periodogram derived from fluctuations in mean daily heart rate. Grouping of participants into four chronotypes is based on weights associated with components derived by nonnegative matrix factorization (overlaid color trace; see Methods), **b)** Mean periodogram for individuals belonging to the same chronotype. Shaded area is ±1 standard deviation, **c)** Example of mean daily heart rate from one individual within each cluster, showing daily average heart rate (in black) smoothed with a 2-, 7-, 9-, and 87-day moving average filter, respectively, and overlaid filtered cycle estimate (in red). Unique scales are given for each example.

Weekly chronotypes were predominately observed in male participants (78.6%, n=55/70), shorter monthly chronotypes were predominantly observed in females (77.3%, n=75/97), longer monthly chronotypes were almost equally observed in males and females (males: 53.5%, n = 84/157), and multi-month chronotypes were predominantly observed in males (62.1%, n=18/29). Despite their similar periods, the shorter monthly (25-day) and longer monthly (35-day) chronotypes appeared to reflect distinct demographic groups due to disparate sex ratios and cross-correlation analysis.

While the cohort-wide multiday chronotypes were identified using wavelet periodogram distributions, multiday heart rate rhythms were identified on an individual level using significant peaks in wavelet periodograms. The distribution of all significant multiday rhythms showed large clusters of individuals with rhythms at approximately 10-weeks, biannual, and annual periods, in addition to clusters associated with weekly and monthly periods (Figure 3). Considering only the single most dominant peak per individual did not change the overall distribution of multiday rhythms (Supplementary Figure 3).

**Figure 3.**
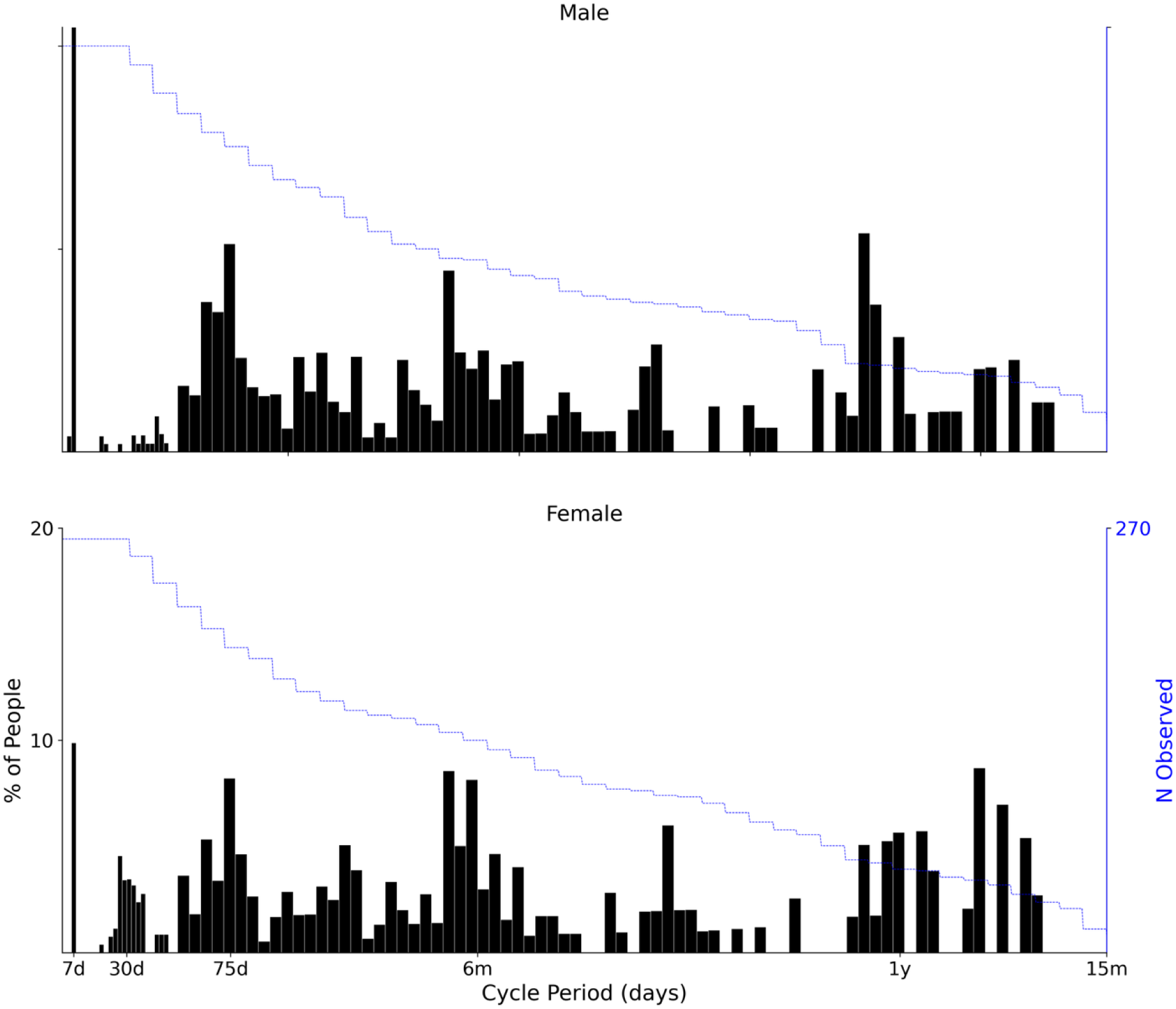
Prevalence of all multiday heart rate cycles. Histograms showing the proportion of people (broken down by sex) with a significant detected rhythm at each cycle period (x-axis), with bar widths increasing from 2 days to 5 days at the 52-day mark. The y-axis on the right-hand side shows the sample size used at each cycle period, noting that the number of people with sufficient data duration decreases with increasing cycle lengths. The maximum period assessed for each individual was one-third of the total data duration, allowing a minimum of three cycles to be observed before significance was determined.

Table 2 shows the number of people who had significant cycles at weekly (between 5-9 days), monthly (between 25-35 days), 10-week (between 8-12 weeks), biannual (between 5-7 months) and annual (between 11-13 months) periodicities. Interestingly, 47.8% of the cohort observed had a significant biannual cycle in their daily heart rate, while 38.9% had a significant annual cycle and 32.8% had a significant 10-week cycle. The most common dominant cycle was biannual (found in 31.9% of people), followed by annual (found in 24.2%) and 10-week (found in 18.9%) cycles. Similar to multiday chronotypes, weekly cycles were predominantly observed in males (68.3% of all observed weekly cycles), and monthly cycles were predominantly observed in females (91.7% of all observed monthly cycles). The mean intra-subject autocorrelation coefficient in the frequency domain over time was 0.69 (average SD = 0.21), demonstrating that these rhythms were relatively stable over time (Supplementary Figures 4 and 5).

**Figure 4.**
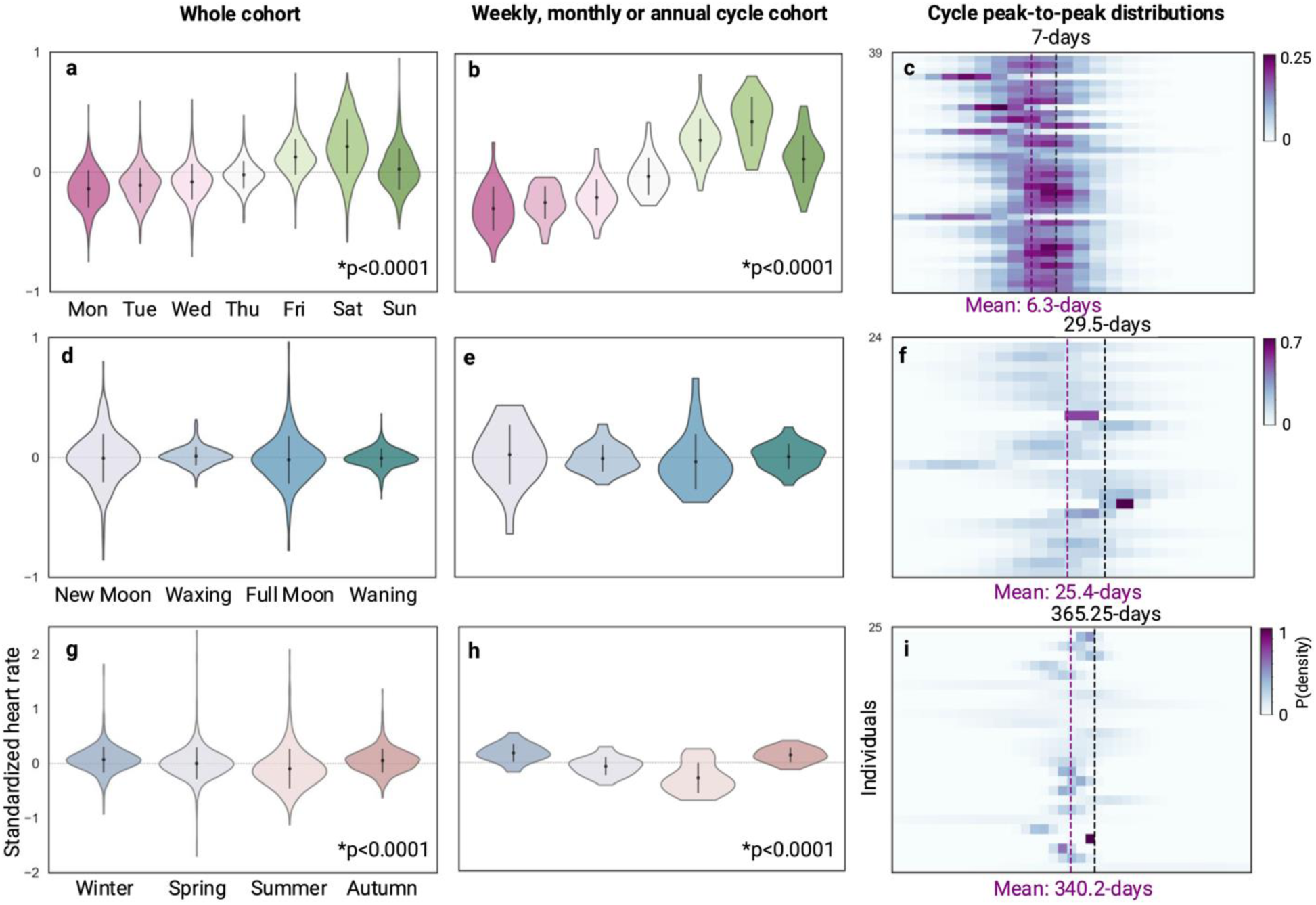
Within-individual standardized heart rate across environmental factors (day-of-week, lunar and seasonal cycles). Day-of-week (7 day) influence demonstrated in **a**, **b** and **c**, lunar (29.5 day) influence demonstrated in **d**, **e** and **f**, and seasonal (365.25 day) influence demonstrated in **g**, **h** and **i**. Heart rate data were first standardized within individuals and then means of the standardized heart rate were calculated per cycle phase (e.g., Winter, Spring, Summer and Autumn), for each individual. The first column (**a**, **d**, **g**) plots the distributions of individuals’ means of standardized heart rate values across phases of the environmental cycle, for the whole cohort with at least 90 days of data (n=558). The second column (**b**, **e**, **h**) also plots the distributions of individuals’ means of standardized heart rate values across phases of the environmental cycle, but only for people whose strongest cycle was (**b**) weekly, defined as 5-9 days, (n=39) (**e**) monthly, defined as 25-35 days, (n=24) or (**h**) annual, defined as 11-13 months (n=25). There was a significant (p<0.0001) difference between at least one comparison of phases in the day-of-week and seasonal conditions, across the whole cohort and strongest cycle cohort, using the ANOVA one-way test. The third column (**c**, **f**, **i**) shows heatmaps of the distributions of peak-to-peak durations (days) for individuals with their strongest cycle being weekly, monthly or annual, respectively. Peak-to-peak durations were calculated using a peak detection algorithm on the filtered cycle, where the passband included frequencies within the full-width-half-maximum of the cycle peak from the individual’s original periodogram. Each row in the heatmap represents one individual. Vertical purple dashed lines represent mean peak-to-peak durations across the cohort, and black dashed lines represent the true environmental cycle duration (7 days, 29.5 days and 365.25 days, respectively). The color intensity represents the probability density (relative likelihood) of observing each peak-to-peak duration in days for each subject. Wider distributions indicate greater variability and reflect non-stationary cycle dynamics, while narrower distributions correspond to more stable, stationary cycles.

**Figure 5.**
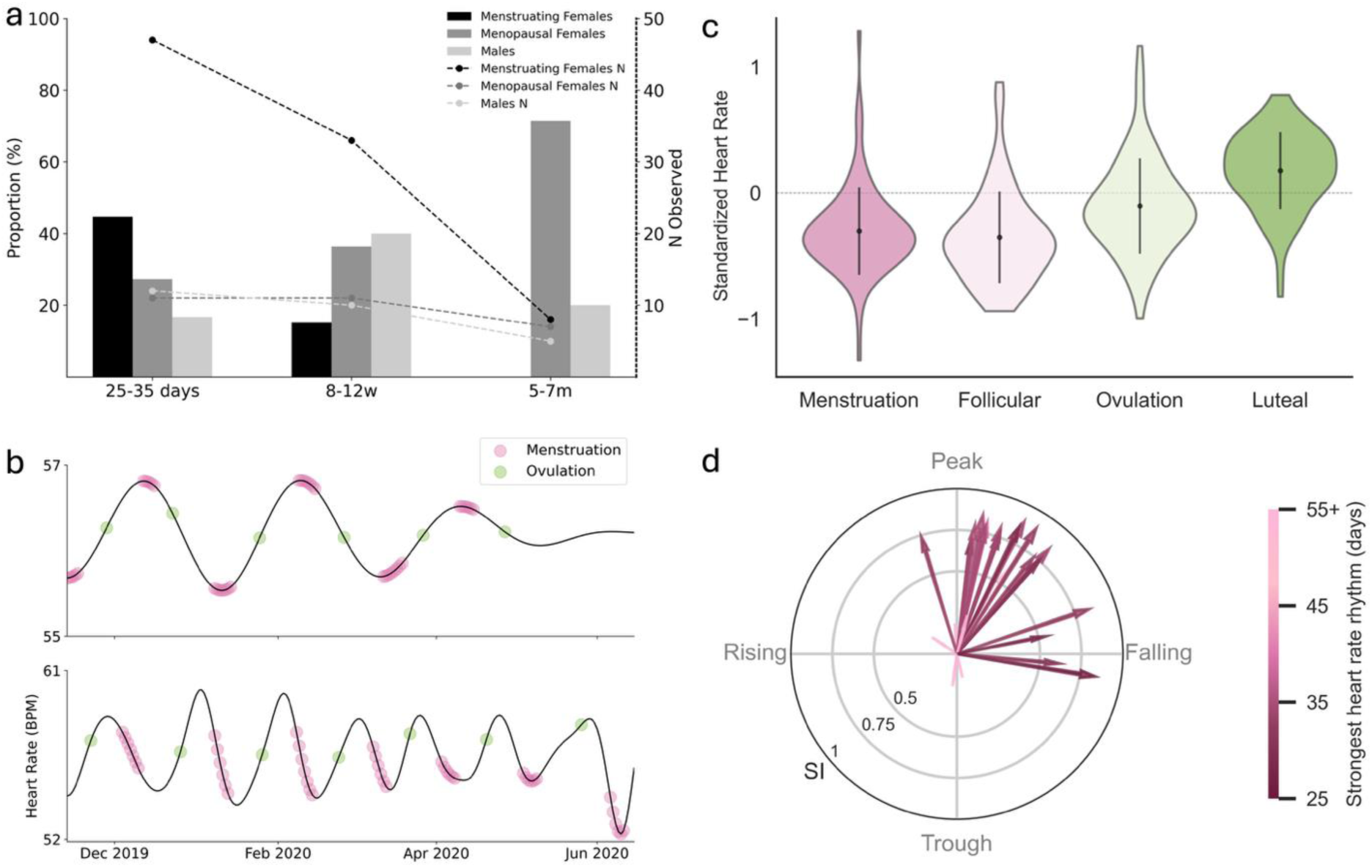
Distribution of multiday heart rate rhythms and association to menstruation within a cohort of 72 adults (49 menstruating females, 11 menopausal females and 12 males). **a.** Prevalence (%) of monthly (25-35 days), 10-week (8-12 weeks) and biannual (5-7 months) cycles in menstruating females (black), menopausal females (dark grey) and males (light grey). The y-axis on the right-hand side shows the sample size used at each cycle period, noting that the number of people with sufficient data duration decreases with increasing cycle lengths. **b.** Two individual menstruating female examples of a heart rate rhythm, i.e., filtered cycle estimate of the individual’s strongest cycle (black), with menstruation and ovulation days dotted in pink and green, respectively. **c.** Standardized daily resting heart rate values across menstrual cycle phases (i.e., menstruation, follicular, ovulation, luteal) in menstruating females, where the ovulation phase was defined as the fertile window (3-days prior to ovulation, ovulation day and 1-day after ovulation). In some cases, when ovulation day was not recorded, ovulation was estimated as the 5-day window before the luteal phase (i.e., the 14-day period before menstruation). Resting heart rate data were first standardized within individuals and then means of the standardized resting heart rate were calculated per menstrual cycle phase for each individual. There was a significant (p<0.0001) difference between at least one comparison of menstruation cycle phases across the cohort using the ANOVA one-way test. **d.** Individual’s Synchronization Index (SI, length of arrows, between 0 and 1) and mean phase of menstruating days with respect to the individual’s strongest heart rate cycle (in days, with darker colours representing shorter rhythms and lighter colours representing longer rhythms), representing the synchrony strength between heart rate cycle phase and menstruation. Only individuals with 5 or more recorded menstruation periods (n=28) were included in the plot; monthly rhythms were the most dominant heart rate rhythms in 61% (n=17) of these females.

**Table 2.** Number of individuals with cycles at different multiday periodicities. Table reporting the breakdowns by sex for individuals whose strongest cycle and significant cycle(s) fall within each cycle-duration category (i.e., 5-9 days, 25-35 days, 8-12 weeks, 5-7 months, 11-13 months). For each category, the overall sample size is provided, reflecting the reduction in sample size for longer periodicities due to variation in length of data recordings across participants.

| Cycle Duration | Sample Size | N Strongest Cycle | % Female | % Male | N Significant Cycle | % Female | % Male |
| --- | --- | --- | --- | --- | --- | --- | --- |
| **Weekly**<br>5-9 days | 523 | 37 (7.1%) | 32.4 | 67.6 | 82 (15.7%) | 31.7 | 68.3 |
| **Monthly**<br>25-35 days | 498 | 25 (5.0%) | 84 | 16 | 48 (9.6%) | 91.7 | 8.3 |
| **10-week**<br>8-12 weeks | 369 | 70 (18.9%) | 42.9 | 57.1 | 121 (32.8%) | 43.8 | 56.2 |
| **Biannual**<br>5-7 months | 226 | 72 (31.9%) | 59.7 | 40.3 | 108 (47.8%) | 52.8 | 47.2 |
| **Annual**<br>11-13 months | 95 | 23 (24.2%) | 30.4 | 69.6 | 37 (38.9%) | 45.9 | 54.1 |

### Environmental effects

Environmental drivers were considered, including day-of-week, lunar phase and seasons. Both the day-of-week and the season had significant (p < 0.0001) effects on heart rate across the cohort, as displayed Figure 4. There was a clear “weekend effect”, with higher heart rate values across the cohort occurring on Friday and Saturday (Figure 4a). There were also weak seasonal effects on heart rate across the cohort, with lower average heart rate during the summer and spring, in comparison to winter and autumn (Figure 4g). Conversely, the lunar cycle did not appear to significantly affect heart rate.

Day-of-week and seasonal effects observed across the cohort were stronger in individuals with significant dominant cycle in the weekly (5-9 day) and annual (11-13 month) range (Figure 4b and 4h), suggesting that these cohort-wide weekly and annual trends are partially driven by a subset of individuals. However, even in these individuals, the peak-to-peak durations of the significant dominant weekly or annual cycle was not closely aligned with the environmental driver (day-of-week or season), showing average peak-to-peak distances of 6.3 days and 340.2 days (rather than the expected 7 days and 365.25 days) and very few individuals exhibiting tight distributions around the fixed-period environmental cycles.

When these three environmental effects were regressed out of individuals’ heart rate signals (Supplementary Figure 6), a marked reduction in the prevalence of the 7-day cycle was observed, while all other cycles remained at a similar prevalence.

### Hormonal effects

The relationship between heart rate rhythms and menstruation was also investigated (Figure 5) in a separate cohort^33^ (Soochow University, China) of 91 menstruating females (mean age 23.52±2.81 years), 12 menopausal females (age=57.08±5.44) and 15 males (age 38.80±10.82 years) who wore Huawei or Fitbit smartwatch devices for at least two months capturing minute-level heart rate data. Within the menstruating females, 84 tracked menstrual periods and 28 additionally recorded ovulation using luteinizing hormone strips. In total across this cohort, 70 had sufficient heart rate data recorded for analysis (47 menstruating females, 11 menopausal females, 12 males) and 75.7% (n=53/70) had a significant multiday heart rate rhythm.

Significant monthly rhythms were most prevalent in menstruating females (observed in 48.9%, n=23/47), followed by menopausal females (27.3%, n=3/11) and males (16.7%, n=2/12); 10-week cycles were most common in males (40%, n=4/10), followed by menopausal females (36.4%, n=4/11) and menstruating females (15.2%, n=5/33); and biannual (5-7 month) rhythms were most common in menopausal females (71.4%, n=5/7) followed by males (20%, n=1/5), with no biannual rhythms observed in menstruating females. No weekly (5-9 day) rhythms were observed in this cohort, and there was insufficient data to observe yearly/seasonal rhythms.

Across the cohort of menstruating females, the menstrual cycle was significantly (p < 0.0001) linked to resting heart rate, with resting heart rate peaking during the luteal phase and at its lowest during the follicular phase (Figure 5c). Menstruating days were strongly synchronized to individuals’ dominant heart rate rhythms in most menstruating females (n=28 females with ³5 recorded menstrual cycles, median synchronization index: 0.75, IQR: 0.60, Range: 0.05-0.90), with menstruating days most often occurring just after the peak of the heart rate rhythm (Figure 5d). The high synchronization index values suggest a coupling of multiday resting heart rate rhythms to circulating reproductive hormones in most menstruating females.

### Demographic and behavioral effects

Figure 6 shows the demographic and behavioral characteristics of each multiday chronotype (weekly, multi-weekly, monthly, multi-month), according to individuals’ sex, circadian chronotype preference (morning/intermediate/evening), average activity level (less, moderately and very active) and average sleep duration (short, <8 hours, and long, ≥8 hours). Activity levels and sleep duration were categorized according to average American guidelines (Supplementary Appendix 6).

**Figure 6.**
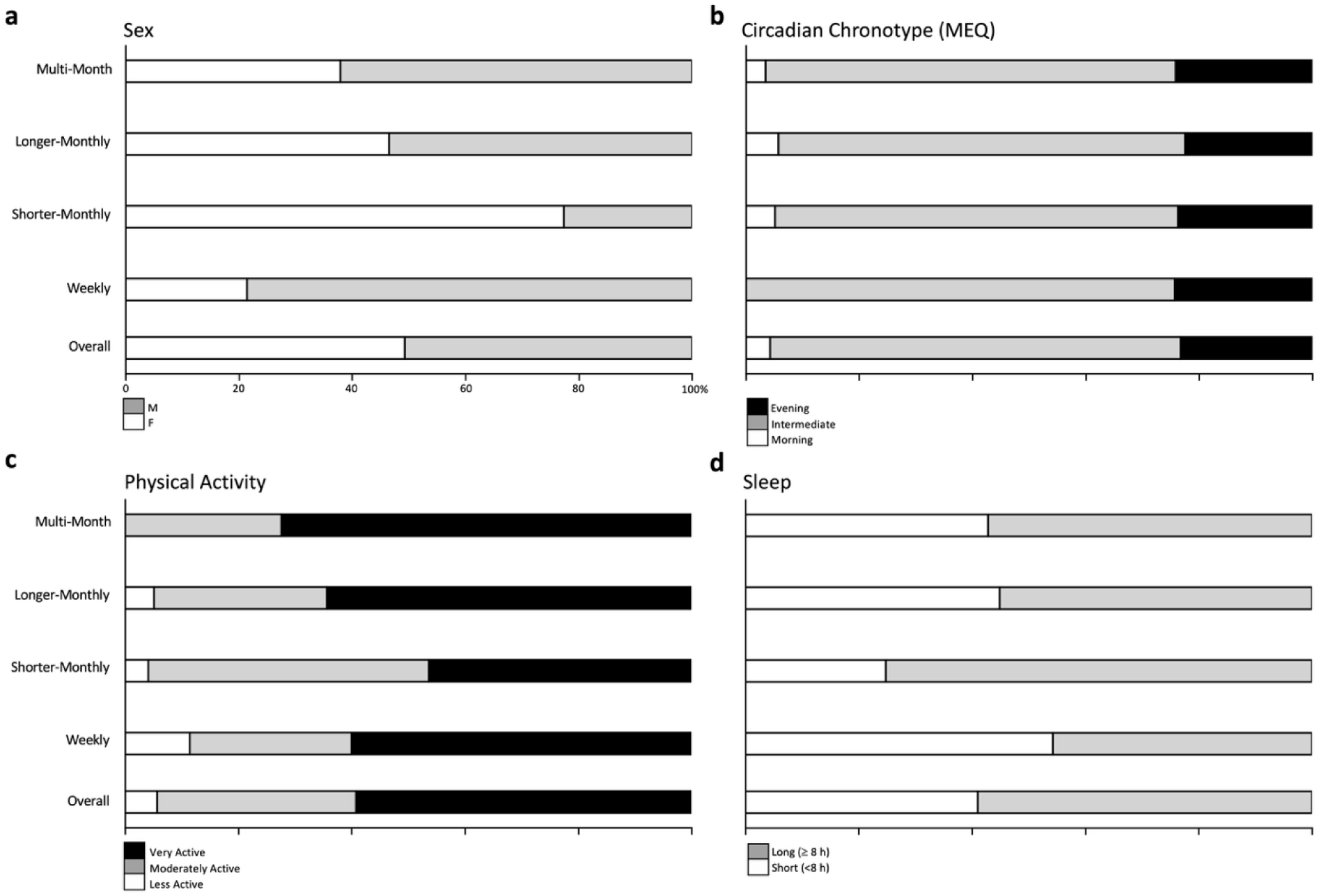
Distribution of multiday chronotypes according to demographic and behavioral factors. Subpanels show the proportion of people with each multiday chronotype based on **a) Sex**, **b) Circadian chronotype**, i.e. the preferred sleep timing assessed from self-reported MEQ survey instrument (here ‘definite’ and ‘moderate’ morning/evening types were combined). **c) Weekly activity level** based on the mean active minutes per week (measured from the wearable device) and defining less active as <150mins per week, moderately active as 150 – 300mins per week and ‘very active’ as >300 mins per week. The distribution of activity minutes is given in Supplementary Figure 7. **d) Sleep duration** recorded from wearable devices, with ‘short sleep’ defined as <8 hours, ‘long sleep’ defined as ³8 hours.

Multinomial logistic regression was used to model significant predictor variables to investigate the associations between these factors and multiday chronotype. As seen in Figure 6 and confirmed by the model (Supplementary Table 2), sex was the strongest predictor of multiday chronotype, with females significantly more likely to have monthly (both shorter- and longer-monthly) multiday chronotypes. Less active individuals showed a similar and significant pattern, with activity levels likely related to sex in this cohort. There were no morning-types in the weekly group, and people with longer sleep were over-represented in the shorter monthly group; however, these trends were not significant.

Finally, the phenomenon of physiological synchrony was explored by comparing the wavelet periodogram distributions and peaks between pairs of individuals with close (i.e., roommates, siblings or friends) and socially distant (i.e., strangers) relationships (Supplementary Figure 8). Close pairs demonstrated significantly higher similarity in wavelet periodogram distributions and peak features than socially distant pairs (p = 0.004).

## Discussion

Infradian, or multiday, rhythms of average heart rate were found in over two-thirds of healthy young (college-aged) adults, commonly with weekly, monthly or longer (e.g., 6-and 12-month) periods. Cycles were not necessarily linked to environmental cues, such as the day-of-week, lunar phase or seasonal transitions. Despite evidence of weekday effects, weekly cycles were not the strongest cycle for most people when environmental effects were regressed out of the model. Indeed, the most common rhythms were biannual (6-month) and 10 weekly cycles, which lack any clear environmental or behavioural explanation. Sex had a significant effect on cycle period, with males being more likely to have weekly and multi-month rhythms, and females more likely to have shorter-monthly rhythms. Both males and females were equally likely to have a longer-monthly chronotype. Given the prevalence of multiday heart rate rhythms amongst healthy young people, it is vital to understand their underlying mechanisms and possible effects on both healthy function and disease states. This study represents a fundamental step towards elucidating widespread, individual-specific infradian rhythms affecting human physiology.

### Prevalence of heart rate rhythms

The widespread detection of multiday rhythms and their heterogenous timescales supports the existence of intrinsic mechanisms alongside known environmental modulation. While to the author’s knowledge, this is the first study to investigate multiday rhythms of heart rate in the general population, endogenous cycles of cardiac activity have previously been identified in individuals with pacemakers^13^, epilepsy cohorts^12,29^, and premature infants^3^. The current work confirms the findings of these earlier studies and further extends the understanding of multiday autonomic rhythms beyond narrow cohorts and disease-specific effects to fundamental physiology. Robust annual and biannual rhythms in heart rate were observed in nearly half of the cohort, consistent with previous observations in clinical populations^13^. While seasonal factors significantly affected heart rate – with peaks in winter and troughs in summer – this modulation likely reflects a complex interplay of environmental photoperiods, temperature, and seasonal shifts in physical activity. Such circannual variation is well-documented in metabolic^34^ and hormonal secretion^9,35,36^. Nevertheless, several lines of evidence argue against a purely exogenous driver for these cycles. First, the peak-to-peak durations of annual heart rate rhythms were not precisely locked to the 365-day solar year, suggesting that the environmental cycle may entrain, but does not entirely account for these oscillations. Second, the high prevalence of biannual (47.8%) and 10-week (32.8%) rhythms cannot be explained by known geophysical or social influences. While biannual rhythms may represent a harmonic of the annual cycle, the 10-week periodicity lacks any clear intrinsic or extrinsic precedent, though it may emulate the quarterly (3-month) cycles observed in patients with cardiac pacemakers^13^. Thus, the evidence in this study points toward a conserved, long-term regulatory mechanism of the autonomic nervous system. Further investigation is required to elucidate the underlying drivers of these biannual and 10-week oscillations, which appear to be fundamental, yet under-recognized, features of human physiology.

Another central finding of this work was the presence of monthly rhythms across the entire cohort, including in males and menopausal females. Menstruation times were strongly synchronized to heart rate rhythm phase in most menstruating females, suggesting a coupling between the autonomic nervous system and circulating reproductive hormones. While previous research has identified links between the menstrual cycle and markers of the autonomic nervous system (including heart rate, temperature, respiratory rates and blood pressure^33^), the presented results extend these findings by suggesting the presence of continuous and robust autonomic fluctuations that can co-oscillate, or synchronize, with reproductive hormones. However, the existence of monthly multiday rhythms in males and menopausal females across this cohort and others^12,24,26^, as well as the prevalence of rhythms at other periodicities, indicates that monthly heart rate rhythms are not solely driven by female reproductive hormones. Indeed, much like the circadian rhythm is primed to entrain to the daily light-dark cycle, it is possible that intrinsic multiday rhythms tend to synchronize to strong biological and environmental drivers.

Nevertheless, the striking differences in multiday rhythm periods and multiday chorotypes between males, menstruating females and menopausal females, suggests that the architecture of multiday coupling changes across sex and life stages. For instance, monthly rhythms were rare in a cohort of 336 individuals with pacemakers^13^, which likely reflects an older cohort; in fact, annual rhythms were by far the most common, found in two-thirds of the cohort, followed by quarterly (12%) and weekly (12%) rhythms. In the present study, weekly rhythms were observed in 15.7% of individuals, predominately males, which may be partially explained by shorter reproductive hormone rhythms documented in males^4^. On the other hand, day-of-week effects clearly influenced weekly rhythms in this cohort, which could be related to weekly behavioral changes, such as higher alcohol consumption or deviations from normal bed- and wake-times over the weekend, both known to transiently elevate resting heart rate^30^. Notably, correcting for day-of-week substantially reduced the prevalence of precise 7-day rhythms, albeit not other about-weekly rhythms. This aligns with previous research in people with epilepsy, where about-weekly rhythms in seizure timing^16^, brain excitability^26^ and heart rate^12^ are frequent but rarely precisely 7 days.

### Physiological mechanisms of heart rate rhythms

There are several proposed theories of the origin and biological processes responsible for controlling and synchronizing biological rhythms in the body, although the mechanisms of multiday rhythms remain unknown. The suprachiasmatic nucleus (SCN) in the hypothalamus is well-recognized as the master circadian clock, controlling daily physiological processes and enabling alignment to environmental conditions^37^. Peripheral clocks are circadian oscillators located in tissue and organs; while they were historically thought to be synchronized by the SCN, emerging evidence suggests that they are not entirely reliant on the SCN^37,38^ and have the ability to synchronize with other organs and tissues through intracellular coupling and paracrine signaling^37^. This ability of peripheral clocks to oscillate independently from the SCN forms the basis for proposed hypotheses as to the origin of multiday rhythms; more specifically, since peripheral clocks are not necessarily entrained to environmental cues (i.e., the light-dark cycle of the solar day) they could oscillate with slower frequencies and follow internal cues^38^. Thus, a possible mechanism of multiday rhythms is that physiological systems consisting of coupled circadian oscillators drive longer, emergent rhythmic dynamics, within a broad range of timescales^39^, consistent with the observed heterogeneity of heart rate cycles.

Beyond individual biological pacemakers, our findings suggest that multiday rhythms are modulated by social and environmental proximity. Individuals in close relationships (i.e., roommates, siblings and friends) exhibited significantly higher similarity in their multiday chronotypes and rhythm periodicities compared to strangers, suggesting a form of interpersonal physiological synchrony – a phenomenon previously documented in short-term contexts such as parent-child interactions^40^, friendships^41^, and those performing group tasks^32,42,43^ and sharing experiences^41^. Our results extend these observations to a much longer timescale, suggesting that shared social zeitgebers may entrain the autonomic nervous system over weeks and months. While anecdotal evidence for long-term synchronization (most notably the ‘McClintock effect’ in menstrual cycles^44^) remains a subject of scientific debate, our data supports the existence of shared multiday rhythmicity among socially connected individuals and, combined with the present findings on menstrual cycle/heart rate rhythm synchronicity, may even partially explain the McClintock effect. While sibling sample sizes were insufficient to suggest a genetic driver in this work, future studies are warranted to decouple the relative contributions of genetic architecture vs shared environmental entrainment in governing these long-term biological clocks.

### Potential implications for health

The characterization of multiday rhythms offers a transformative framework for individualized, chronotherapeutic disease management. This approach is validated in the management of chronic neurologic conditions like epilepsy, where monitoring multiday seizure cycles can improve health outcomes and quality-of-life^24,45,46^. The striking similarity in prevalence between the heart rate rhythms observed here in healthy adults and in clinical cohorts^13,15,24^ suggests that multiday periodicity is a fundamental property of the human nervous system. While the biological impact of multiday rhythms is unexplored, recent research suggests links to salivary cortisol^47^ and brain excitability^12,48^. Thus, monitoring these rhythms could provide a window into periods of heightened susceptibility to stress, potentially transforming the management of psychiatric disorders that fluctuate over extended timescales, such as seasonal affective disorder and bipolar disorder. Moreover, multiday heart rate rhythms have been found in cardiovascular disease, alongside exacerbations in disease symptoms at specific circadian^49^, seasonal^19^ and even weekday^50^ phases. While the cyclic risk profiles of cardiac disease have mostly been studied at the population-level, tracking individual-level cycles could be an important time-varying biomarker of the risk of cardiac arrythmias, or even heart attacks.

In women’s health, routine tracking of infradian biology has already seen widespread adoption, with tracking of hormonal and menstrual cycles used for fertility and lifestyle planning^51^. However, there are limitations in accuracy and consideration of variability between individuals, especially when the cycle does not conform to a 28-day timeframe^51^. The findings presented here suggest that heart rate rhythms may be utilized to improve hormonal and menstrual cycle tracking, particularly during periods of heightened stress^52^, which is known to affect menstrual period regularity^52^. Thus, by leveraging the acquisition of heart rate data via wearable sensors, heart rate rhythms have potential to provide a longitudinal proxy for reproductive and stress hormones, although further study is undoubtedly required. Nonetheless, this shift toward continuous biometric monitoring reduces reliance on discrete manual logging, offering a scalable approach to characterize individual variability and non-stationarity in reproductive physiology across diverse populations.

### Limitations and future work

This study primarily analyzed the NetHealth dataset, which comprised daily averaged heart rate from a relatively homogeneous cohort of young adults aged 17–23 years. Although participants varied in demographic background and had diverse geographic origins across the United States, they shared broadly similar lifestyles as students at a single university and resided predominantly in the same location during the four-year study period, which may limit generalizability. Future work should test the robustness of these multiday rhythms in larger, more diverse cohorts spanning wider age ranges, geographic contexts and socioeconomic backgrounds, and should incorporate family- or twin-based designs to quantify potential genetic contributions.

A second limitation is the temporal resolution of the wearable data used in this work. Future studies should aim to analyze a higher resolution dataset with minute-by-minute or hourly heart rate values to obtain reliable measurements of resting heart rate. While results from our second, higher-resolution cohort (menstrual cohort) are consistent with the NetHealth results, obtaining accurate resting heart rate measurements may remove uncertainty in environmental and behavioral influence that we were unable to entirely account for in this work. Finally, detecting free-running rhythms of unknown period in noisy, non-stationary real-world data remains methodologically challenging, and there is no accepted gold-standard pipeline. Continued development of validated methods will be required to improve accuracy and to characterize within-individual cycle variability – an essential consideration for population-scale studies of multiday physiology.

## Conclusion

This study provides the first large-scale characterization of infradian (multiday) heart rate rhythms in a healthy population, demonstrating that human cardiac physiology is governed by a complex, multi-layered temporal architecture. The findings revealed that two-thirds of individuals exhibit robust multiday cycles, often independent of exogenous environmental influences. By identifying distinct multiday chronotypes associated with sex, sleep, and social proximity, we establish these oscillations as fundamental features of individual autonomic regulation. The integration of these rhythms into wearable health platforms offers an increasingly relevant framework for improving women’s health tracking, monitoring and managing fluctuating neurological and psychiatric conditions, and identifying abnormalities in cardiovascular health. Ultimately, these results reveal an under-recognized dimension of human physiology, reinforcing the need to investigate these rhythms over larger, more diverse cohorts and in the context of disease and personalized medicine.

## Methods

### Dataset and preprocessing

This study analyzed data from the University of Notre Dame’s NetHealth project (Indiana, U.S.A.), a publicly available longitudinal dataset that followed a cohort of university students over four years (August 2015–May 2019). Participants (n=623) wore Fitbit smartwatches (either Charge HR or Charge 2 model), which continuously recorded health-related metrics, including daily heart rate, physical activity and sleep metrics^30^. The individual wearable device data included timestamped daily mean and standard deviation of heart rate recorded from the smartwatch (in beats per minute), as well as minutes of daily physical activity (defined by Fitbit metabolic equivalent tasks (MET) model, i.e., 1.5-3.0 METs categorized as ‘lightly active minutes’, 3.0-6.0 METs for ‘fairly active minutes’, and >6.0 METs for ‘very active minutes’) and daily sleep metrics (including estimated sleep time, wake time and duration).

In addition to the wearable data, this study incorporated selected survey measures and demographic data available within the NetHealth dataset. Demographic data included age and sex. Surveys included the Morningness–Eveningness Questionnaire (MEQ), which assessed self-reported circadian preference or chronotype, and a Social Network survey, in which individuals in the cohort reported the 20 other individuals they spent the most time with over the previous few months (often containing other participants in the study). A summary of the participant demographics and characteristics can be found in Supplementary Table 1. The full dataset and accompanying documentation are accessible at https://sites.nd.edu/nethealth/.

This study used individual daily heart rate data to identify multiday rhythms. Participants with at least 3 months of recorded heart rate were included in the analysis. To account for the well-known relationship between heart rate and physical exertion, daily physical activity was regressed out of the daily heart rate to obtain a detrended heart rate signal for each participant^29^. A standard linear model was applied:

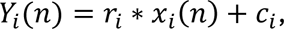

Where 𝑌_𝑖_(𝑛) represents the mean heart rate for a given day, *n*, 𝑥_𝑖_(𝑛) is the number of active minutes per day (taken as the sum of moderate and high intensity activity minutes defined by Fitbit as ‘fairly active’ (3-6 METs) and ‘very active’ (>6METs), respectively) and 𝑐_𝑖_ and 𝑖 are the model correlation coefficient and intercepts for a given participant. The model excluded days when the number of active minutes was zero and was fitted using ordinary least squares linear regression to minimize the residual sum of squares between the observed data and the linear approximation. Thus, daily mean heart rate was approximated as 𝑌_𝑖_(𝑛) with the daily physical activity component (𝑟_𝑖_ ∗ 𝑥_𝑖_(𝑛)) subtracted.

Spectral analysis was conducted on the highest-quality data segment, defined as the longest continuous segment (of at least 3 months) with fewer than 20% missing data points. Missing segments were interpolated with the global average (validated in previous work^12^), provided that the total missing data duration was not greater than 20% of the individual’s segment duration. The longest continuous segment of detrended heart rate data was z-standardized (by subtracting the mean and dividing by the standard deviation) prior to wavelet analysis.

### Multiday rhythm estimation

The pre-processed, detrended heart rate data was analyzed using a wavelet decomposition method with a Morlet wavelet transform, similar to previous approaches for detecting free-running multiday rhythms^12,15^. Resulting spectrograms were averaged across time to generate the global wavelet power or periodogram. Significant multiday cycles were identified as peaks in the wavelet periodogram that crossed the global wavelet 95% significance threshold (α>0.05). Candidate cycle periods ranged from 4 days to a maximum of one-third of an individual’s recording length (i.e., a minimum of three cycles had to be observed) with a step size of 12 hours. Restricting the range of periods analyzed helped reduce boundary (edge) effects in the wavelet transform, consistent with the cone of influence principle.

Each significant peak in the periodogram was defined by its prominence, i.e., the extent to which a peak stands out from the local baseline, measured as the vertical distance between the peak and the lowest contour that surrounds it. When more than one significant peak was detected, peaks were assessed for distinctness. Peaks were considered distinct significant cycles if they fell outside the full-width-half-maximum range of a neighboring, more prominent peak. Thus, individuals’ multiday cycles were defined as all significant, distinct peaks in the periodogram, with the most *dominant* cycle defined as the most prominent peak.

### Multiday chronotypes

The concept of multiday chronotypes has previously been proposed to classify common cycle periods across a population of people. For instance, in neurological cohorts, multiday seizure cycles are commonly observed at about-weekly, about-monthly and seasonal periods^24^. This study used non-negative matrix factorization (NNMF) to quantitatively describe the cyclical trends and clusters across the population without pre-defining set chronotypes. NNMF factorizes a matrix of averaged power values across periods in each individual’s dataset into two non-negative matrices: weights and components. The optimal number of components was chosen by using a cross-correlation metric described in the supplement (Supplementary Figure 2). Thus, for each individual, the strongest weight component in the NNMF can be identified and this allows sorting of individuals into chronotypes based on their dominant heart rate cycles.

### Statistical information

#### Cycle Significance

To determine significant cycles for individuals, a global wavelet significance threshold was applied to the global wavelet periodogram. This threshold was determined through an in-built wavelet significance line in the PyCWT Python package which utilizes wavelet analysis and statistical analysis. A time-average null-hypothesis significance test was implemented with a significance level of 95%. Any peaks surpassing this threshold in the wavelet power periodogram were deemed statistically significant and thus a true underlying rhythm in the data.

#### Environmental Effects

The influence of three environmental variables on multiday cycles was investigated: day of the week (Monday to Sunday), lunar phase (new moon, waning, full moon or waxing) and season (spring, summer, autumn or winter). Daily mean heart rate was standardized (z-scored within individuals) and mapped to environmental cycle phases, and individuals’ mean values were calculated for each environmental cycle phase. Individuals were grouped at cohort levels, and significant differences in standardized heart rate across environmental cycle phases were tested using one-way ANOVA tests.

A generalized additive model was fitted to standardized heart rate to quantify the influence of the three environmental variables. The residuals of the model, i.e. the resulting signal after subtracting the sum of the environmental effects, reflecting variation not explained by these cyclical factors, were then used as inputs for a cohort-level multiday cycles detection analysis.

#### Menstrual cycles

To investigate the influence of the menstrual cycle on heart rate and multiday cycles in heart rate, we analysed longitudinal wearable and menstrual cycle data captured as part of a study by Soochow University, China^33^. This publicly available dataset includes 91 menstruating females (mean age= 23.52±2.81 years), 12 postmenopausal females (age=57.08±5.44) and 15 males (age=38.80±10.82) who wore Huawei or Fitbit smartwatch devices for at least two months capturing minute-level heart rate data. The mean recording duration of this data is 368.46 days (standard deviation SD 252.75 days). Within the menstruating females cohort, 84 tracked menstrual periods and 28 additionally recorded ovulation using luteinizing hormone strips.

Using the minute-level heart rate data, we extracted daily resting heart rate values as the mean of the bottom 30% of minute-by-minute heart rate data per day, thus estimating average heart rate during sleep. Daily resting heart rate data were analyzed in the same way as the daily mean heart rate data from the NetHealth dataset to extract multiday rhythms (details in *Multiday rhythm estimation* section). Only participants with at least 3 months of continuous recorded heart rate (defined as greater than 80% adherence) were included in the analysis.

The relationship between menstrual cycle and heart rate was investigated on a cohort level and an individual level. On a cohort level, daily resting heart rate was standardized (z-scored within individuals) and mapped to menstrual cycle phases (i.e., menstruation, follicular, ovulation, luteal), and individual’s mean resting heart rate values were calculated for each menstrual cycle phase. Significant differences in standardized resting heart rate across menstrual cycle phases were tested using one-way ANOVA tests. On an individual level, the synchronization index (SI) of the menstrual days with respect to phases of the individual’s strongest heart rate cycle (any duration in days) was calculated. The SI is always between 0 and 1 and represents the strength of the synchrony between heart rate cycle phase and menstruation. Only individuals with 5 or more recorded menstruation periods were included in this individual analysis.

#### Multinominal Logistic Regression

Multinomial logistic regression model was applied to the chronotype classifications to explore links between multiday chronotype (as the outcome) and specific demographic or behavioral factors. The log-odd probabilities used a baseline category for multiday chronotype (here, weekly) and for the categorical data (here: male, very active, and longer sleep), providing a coefficient and corresponding p-value for each outcome. Circadian preference was excluded from the model as some categories had a zero-count leading to complete separation. The model coefficients provide an indicator of the change in log-odds of belonging to a given outcome category in comparison to the baseline category (i.e., higher coefficient values reflect greater odds that a given demographic or behavioral category belongs to the outcome chronotype category), similar to effect size. The log-odds were converted into predicted probabilities of the factors being associated with specific chronotypes.

#### Mann-Whitney U Statistical Test

The influence of synchronicity of multiday cycles amongst individuals was considered. Each individual’s global wavelet periodogram was compared against other individuals (who were also study participants) in their self-reported social network. Individuals in the social network were grouped by interpersonal closeness, specifically ‘Close’ and ‘Not Close’ categories (as described in detail in Supplement), and a peak similarity metric was employed to compare periodograms. A nonparametric Mann-Whitney U statistical test was performed to assess if there was a significant difference between the peak similarity values of ‘Close’ individuals and ‘Not Close’ individuals across the cohort.

## Data availability

The datasets analyzed during the current study are available in the University of Notre Dame’s (USA) NetHealth repository, https://sites.nd.edu/nethealth/ (heart rate for main cohort) and Mendeley, https://doi.org/10.17632/v58stpfcnm.1 (heart rate with menstrual cycles timing).

## Acknowledgments

Thank you to Dr. Mark Bennett for his assistance in developing the environmental effects model.

