## Supplement for "Multiday rhythms modulate human heart rate: an observational study in healthy adults"

### Supplementary Material

**Supplementary Table 1. Participant demographics and characteristics**

| | Number of Participants |
|---|---|
| **U.S. Region of Origin** | |
| **New England**<br>***(i.e., Connecticut, Maine, Massachusetts, New Hampshire, Rhode Island, Vermont)*** | 24 |
| **Middle Atlantic**<br>***(i.e., New Jersey, New York, Pennsylvania)*** | 81 |
| **East North Central**<br>***(i.e., Illinois, Indiana, Michigan, Ohio, Wisconsin)*** | 222 |
| **West North Central**<br>***(i.e., Iowa, Kansas, Minnesota, Missouri, Nebraska, South Dakota, North Dakota)*** | 54 |
| **South Atlantic**<br>***(i.e., Delaware, Florida, Georgia, Maryland, South Carolina, North Carolina, Virginia, West Virginia, Washington D.C.)*** | 71 |
| **East South Central**<br>***(i.e., Alabama, Kentucky, Mississippi, Tennessee)*** | 15 |
| **West South Central**<br>***(i.e., Arkansas, Louisiana, Oklahoma, Texas)*** | 34 |
| **Mountain**<br>***(i.e., Arizona, Colorado, Idaho, Montana, Nevada, New Mexico, Utah, Wyoming)*** | 17 |
| **Pacific**<br>***(i.e., Alaska, California, Hawaii, Oregon, Washington)*** | 65 |
| **Other** | 40 |
| **Self-Reported Exercise Frequency Summer 2015** | |
| **3 or more times a week** | 305 |
| **1-2 times a week** | 124 |
| **1-2 times a month** | 30 |
| **Less than 1-2 times a month** | 13 |
| **Not at all** | 4 |
| **(No Response)** | 147 |
| **Self-Reported Physical Activity** | |

| | **Summer 2016** | **Fall 2016** | **Spring 2017** | **Fall 2017** | **Spring 2018** | **Spring 2018** |
|---|---|---|---|---|---|---|

| | | | | | | |
|---|---|---|---|---|---|---|
| **Very Inactive** | 33 | 43 | 44 | 46 | 29 | 17 |
| **Inactive** | 78 | 78 | 84 | 62 | 51 | 25 |
| **Moderately Active** | 47 | 45 | 48 | 47 | 40 | 23 |
| **Active** | 120 | 114 | 94 | 93 | 76 | 48 |
| **Very Active** | 123 | 90 | 91 | 64 | 67 | 73 |
| **(No Response)** | 222 | 253 | 262 | 311 | 360 | 437 |
| **PSQI Chronotypes** | | | | | | |
| | **Fall 2015** | **Spring 2016** | **Summer 2016** | **Fall 2016** | **Spring 2017** | **Fall 2017** |
| **Good Sleep Quality** | 122 | 76 | 123 | 40 | 48 | - |
| **Poor Sleep Quality** | 329 | 452 | 275 | 306 | 291 | - |
| **Severe Sleep Difficulties** | 13 | 17 | 2 | 20 | 17 | - |
| **(No Response)** | 159 | 78 | 223 | 257 | 267 | - |

#### Appendix 1: Prevalence of heart rate cycles

Overall, 48.2% (175/363) of people had a single dominant significant cycle and 34.7% (126/363) exhibited two significant cycles Supplementary Figure 1a. There was a significant drop-off in the proportion of people with over two significant cycles, with around 11.8% exhibiting three cycles and a further reduction of 4.4% exhibiting four cycles. The notable drop-off in the proportion of people exhibiting over two cycles is likely due to the inter-individual variability in dataset lengths, over which longer cycles might not be observed due to these limitations. For individuals with at least two significant cycles, those with a strongest cycle between 60-80days commonly shared a weekly (7-day) cycle as their strongest secondary cycle (Supplementary Figure 1b). Similarly, those with a strongest cycle between 168.5-195days exhibited a strongest secondary cycle between 75.5-86days or between 99-113days.

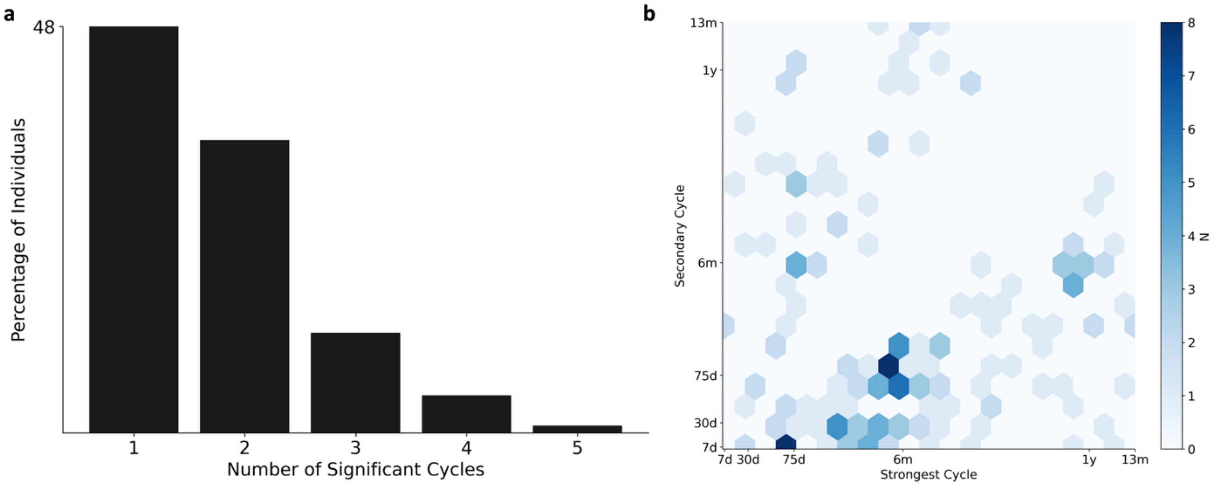


**Supplementary Figure 1: (a) Hist/Distribution of number of cycles. (b) Hexbin visualization of the dominant two cycle periods in people with at least two cycles. Secondary cycle is the second most prominent peak in the wavelet periodogram.**

#### Appendix 2: Selection of Chronotypes with NNMF

The cross-correlation between NNMF components revealed the highest cross-correlation values between features for four components (Supplementary Figure 2). While both 3 and 5 components exhibited close cross-correlation values, neither were chosen due to oversimplification of features when conducting NNMF with 3 components and the lack of significant information added with 5 components. While the NNMF conducted with 5 components revealed the presence of a 'multi-week' chronotype, this chronotype appeared to be an amalgamation of individuals who did not display inter-individual alignment to a specific cycle period; as such, individuals displayed a dominant peak anywhere between 14 – 45 days, thus these were considered to have a 'multi-week' chronotype in the 5 component NNMF iteration. However, since conducting NNMF with either 3 or 4 components resulted in the loss of this 'multi-week' chronotype, and due to the lack of inter-individual alignment to a specified peak, 4 components was deemed appropriate for analysis and breakdown of the chronotypes. This is further substantiated by the increase in the cross-correlation value at 4 components versus both 3 and 5 components in Supplementary Figure 2 below.

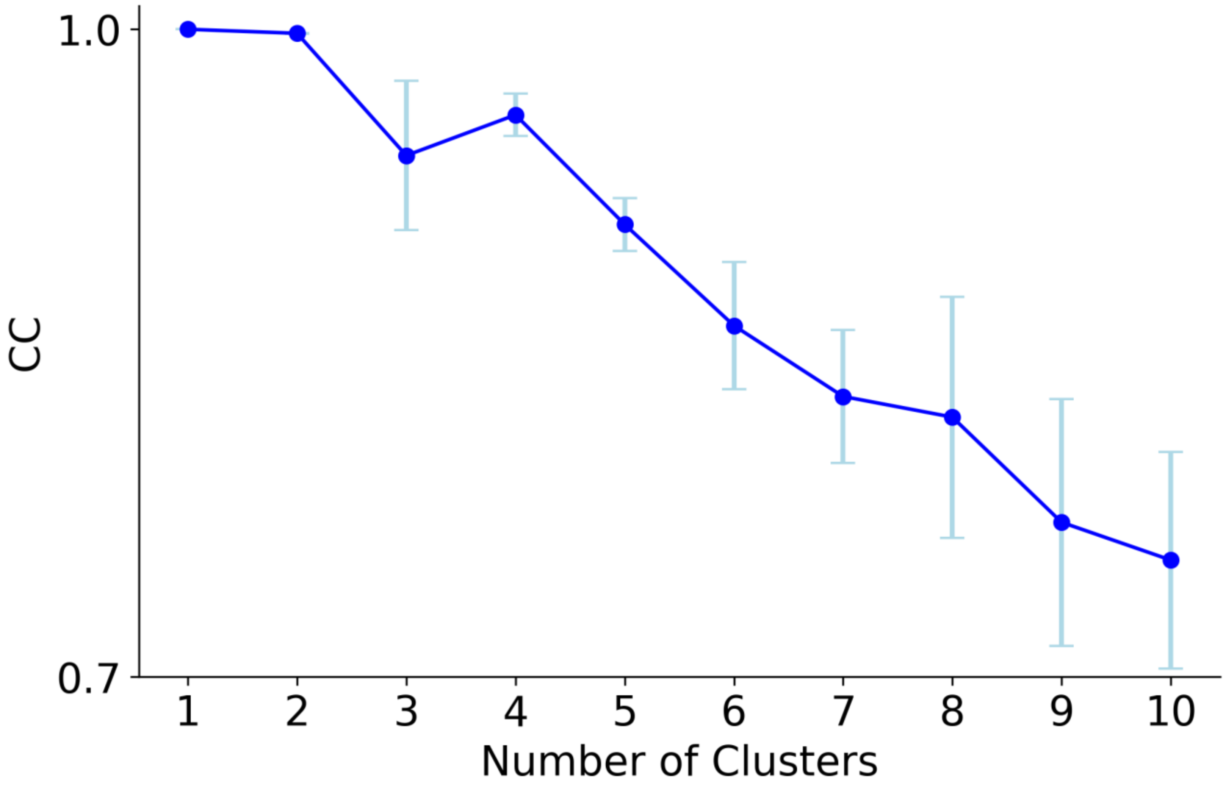


**Supplementary Figure 2: Cross correlation between component groups derived from NNMF decomposition**. The plot displays cross-correlation values between component groups (clusters 1- 10), with each decomposition computed by seeding the algorithm with 10 different initial matrices.

#### Appendix 3: Distribution of multiday heart rate cycles

The distribution of the most dominant multiday heart rate cycle across the population (Supplementary Figure 3) displayed similar clusters when considering all cycles across the population (Figure 3), with peaks in the histogram corresponding to weekly (7-day), monthly (30-day), 75-days, 6-month, and 1-year periods. Compared to the distribution of all cycles across the population (Figure 3), the proportion of people with a dominant shorter cycle (i.e., weekly, monthly and 75-day) appeared to be half the overall distribution whilst the longer cycles (i.e., 6-month cycles and 1 year) although still prominent, remained a similar proportion of the distribution.

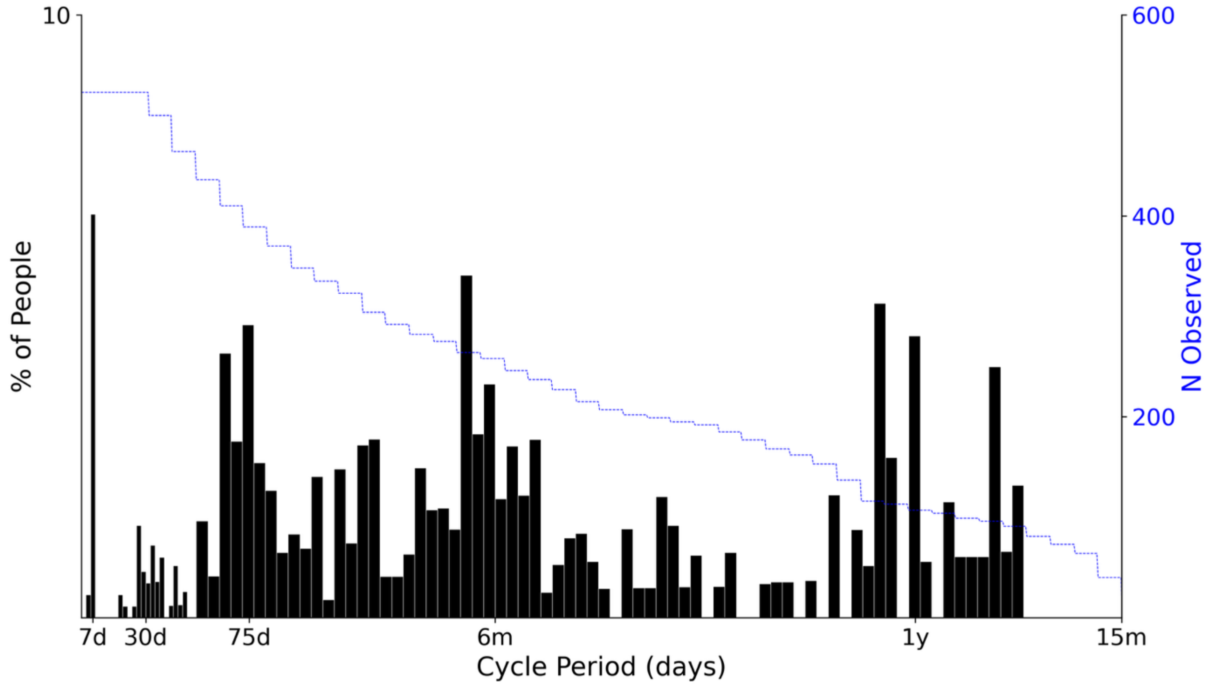


**Supplementary Figure 3: Distribution of the most dominant multiday heart rate cycle amongst individuals.** Histogram showing the proportion of people with a dominant multiday rhythm at each cycle period (x-axis), with bar widths increasing from 2 days to 5 days at the 52-day mark. The y-axis on the right-hand side shows the sample size used at each cycle period.

#### Appendix 4: Stability of multiday rhythms

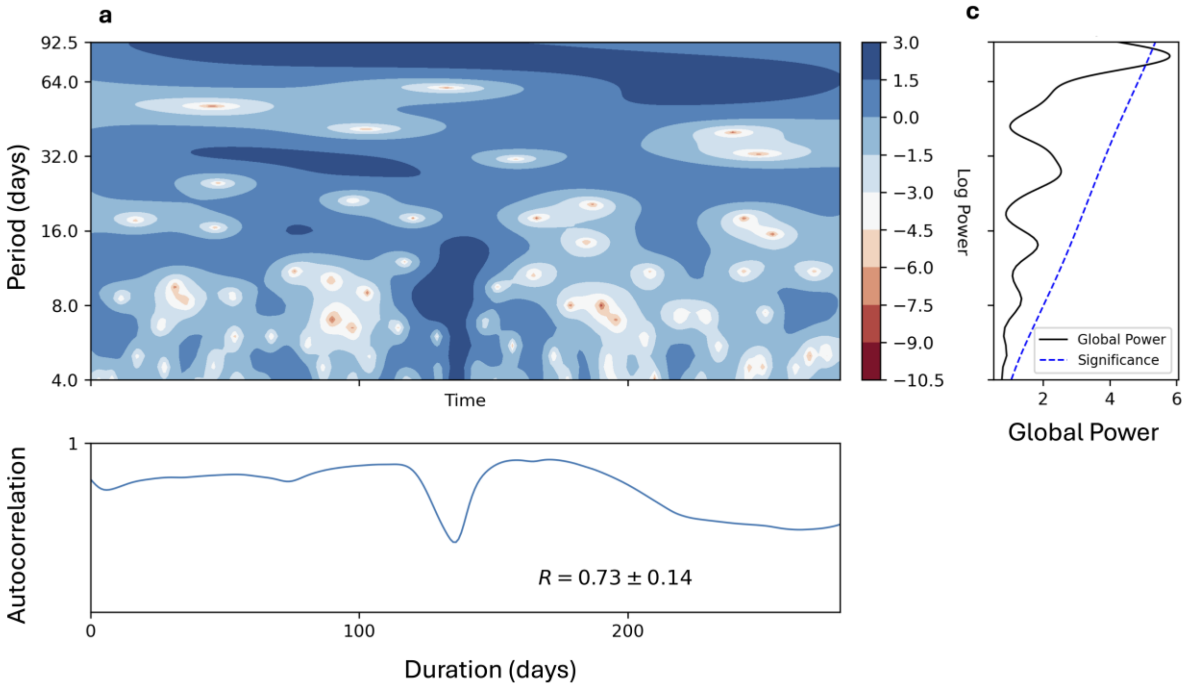


**Supplementary Figure 4:** Stability of multiday rhythms over time using autocorrelation. (a) Spectrogram of one subject over 280 days. (b) Autocorrelation Pearson coefficients between average periodogram (shown in (c)) and spectrogram at each time-point where power was calculated for all frequency bins. (c) Average periodogram revealing one significant peak at approximately 12 weeks.

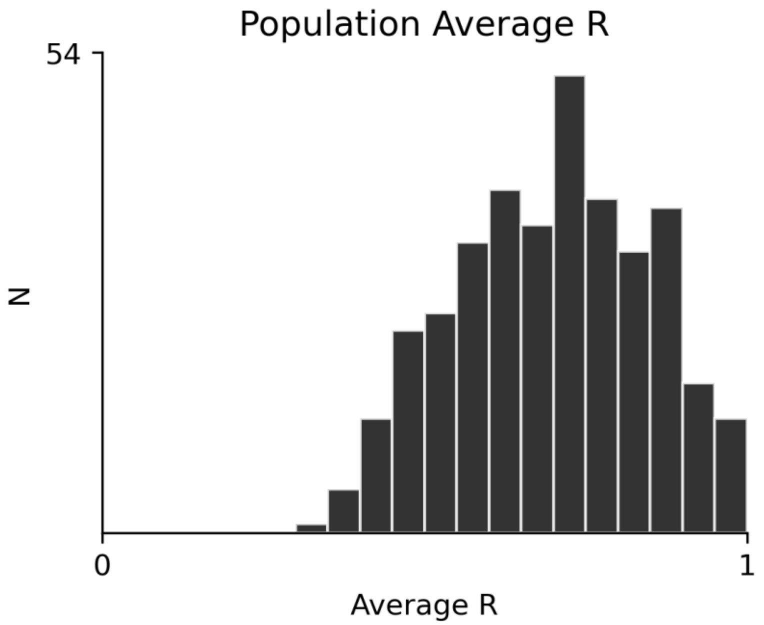


**Supplementary Figure 5:** Distribution of average correlation values (R) across all participants.

#### Appendix 5: Environmental effects

The distribution of all cycles across the cohort was found following the regression of environmental effects (i.e., day-of-week, lunar phase, seasons, and exercise) from the dataset (Supplementary Figure 5). The main striking difference between the distribution prior to and after regression is the diminished weekly 7-day cycle. While the marked 75-day rhythm appears more visually prominent, this cycle and all other cycles remained at a similar proportion of prevalence in the population.

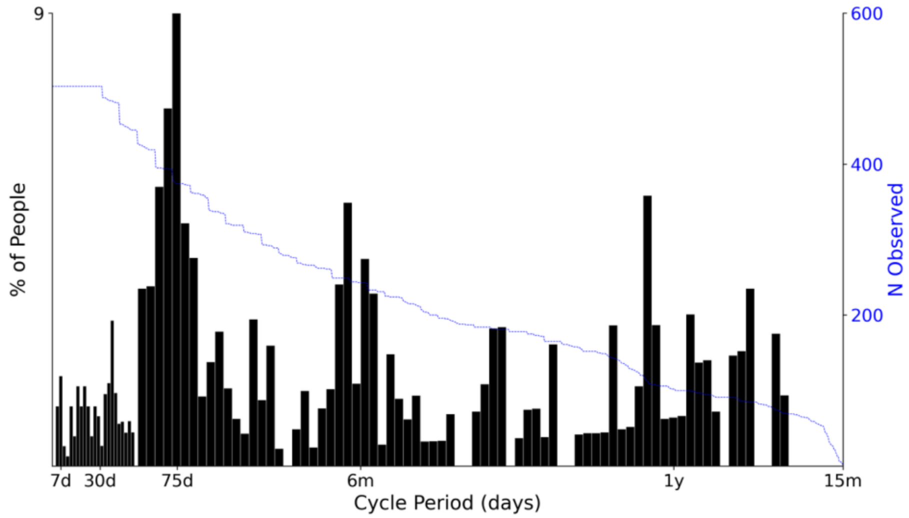


**Supplementary Figure 6. Histogram of all cycles across the cohort after regressing out all environmental effects (i.e., day-of-week, lunar phase, seasons, and exercise).** Histogram showing the proportion of people with a dominant multiday rhythm at each cycle period (x-axis), with bar widths increasing from 2 days to 5 days at the 52-day mark. The y-axis on the right-hand side shows the sample size used at each cycle period.

#### Appendix 6: Exercise and Sleep Distributions

The distribution of weekly exercise and sleep across the cohort was determined in order to generate the respective subplots in Supplementary Figure 6. To analyze the relationship between these two factors and the chronotypes, individuals in the cohort were separated into defined categories for exercise and sleep.

The average American guidelines for weekly exercise of 150-300 minutes (i.e., 2.5 to 5 hours) per week, although a National Health Interview Survey found that only 24.2% of American adults met these guidelines. For this cohort, it seemed that most either met the guidelines or exceeded them (as seen in Supplementary Figure 6a), suggesting these guidelines were optimal criteria to categorize weekly physical activity. Using these guidelines, three distinct categories were delineated to classify varying levels of physical activity: 'Very Active', 'Moderately Active', and 'Less Active'. Thus, individuals' weekly physical activity levels were designated as 'less active' for less than 2.5 hours, 'moderately active' as between 2.5-5 hours, and 'very active' as more than 5 hours per week.

The sleep duration distribution (Supplementary Figure 6b) lay around an average of 8 hours of sleep each night, which aligned with the National Sleep Foundation recommended 7-9 hours of sleep for young adults and is consistent with literature; thus, sleep categories for this cohort were defined as less than 8 hours for 'short sleep' and more than 8 hours for "long sleep".

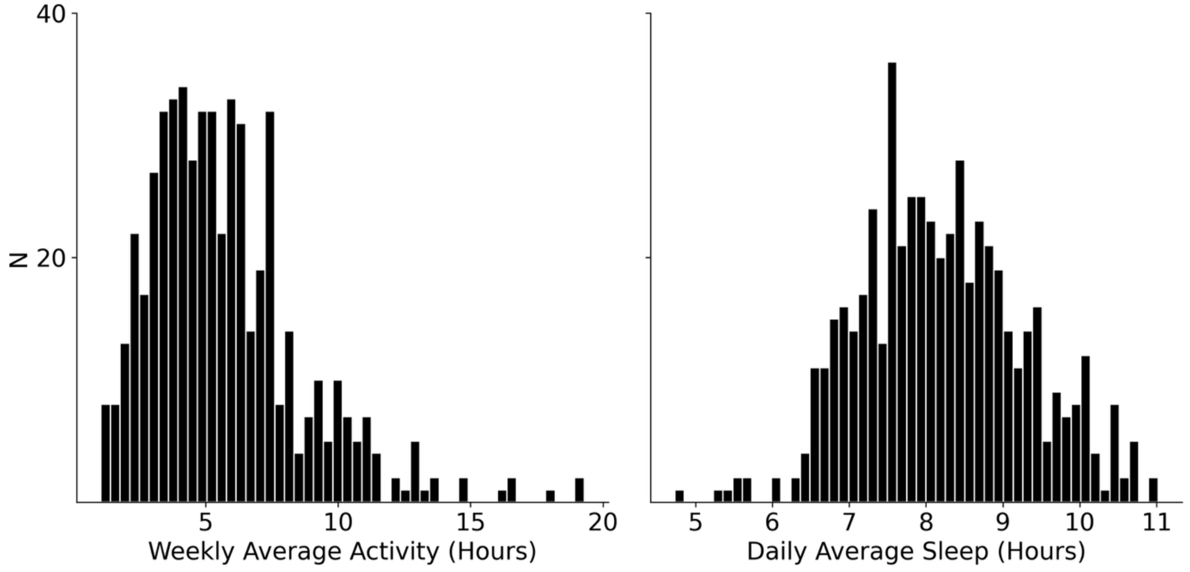


**Supplementary Figure 7. Distribution of weekly activity and daily sleep duration across the cohort.** Figure displays the weekly average activity (hours) on the left and the daily average sleep (hours) on the right. (a) The average weekly activity was computed as the sum of activity minutes where the activity level was ≥3.0 METs. (b) Daily sleep averages were calculated for individuals with at least 14 days of sleep data, where sleep was defined as ≥4 consecutive hours, allowing interruptions of less than one hour.

#### Appendix 7: Links between chronotypes and demographics/behavioral factors

A multinomial logistic regression model was utilized to explore links between multiday chronotypes and demographics or behavioral factors. The outcomes of this model are shown in Supplementary Table 2.

**Supplementary Table 2: Multinomial logistic regression outcomes**. Holding variables constant (i.e., weekly chronotype, Very Active' activity levels, Male, and Longer Sleep), a multinomial logistic regression model is fit to the data to predict an individual's chronotype category from their activity level, sex, and sleep duration. Each of these categories is listed under the 'term' column next to the various chronotypes (y-level column). There are six corresponding columns that describe an estimate (regression coefficient), standard error, statistic (z-value), p-value, as well as low and high confidence levels (95% confidence interval).

| y.level | term | estimate | std.error | statistic | p.value | conf.low | conf.high |
|---|---|---|---|---|---|---|---|
| Longer-Monthly | (Intercept) | 0.551 | 0.273 | 2.020 | 0.043 | 0.016 | 1.086 |
| Longer-Monthly | activityModerately Active | -0.349 | 0.343 | -1.016 | 0.310 | -1.021 | 0.324 |
| Longer-Monthly | activityLess Active | -1.985 | 0.644 | -3.080 | 0.002 | -3.248 | -0.722 |
| Longer-Monthly | sexFemale | 1.496 | 0.393 | 3.805 | 0.000 | 0.725 | 2.266 |
| Longer-Monthly | sleepShorter Sleep | 0.011 | 0.312 | 0.037 | 0.971 | -0.600 | 0.623 |
| Shorter-Monthly | (Intercept) | -0.624 | 0.343 | -1.820 | 0.069 | -1.296 | 0.048 |
| Shorter-Monthly | activityModerately Active | 0.187 | 0.384 | 0.488 | 0.626 | -0.565 | 0.940 |
| Shorter-Monthly | activityLess Active | -1.980 | 0.739 | -2.679 | 0.007 | -3.429 | -0.532 |
| Shorter-Monthly | sexFemale | 2.570 | 0.437 | 5.876 | 0.000 | 1.713 | 3.427 |
| Shorter-Monthly | sleepShorter Sleep | -0.551 | 0.378 | -1.459 | 0.144 | -1.292 | 0.189 |
| Multi-Month | (Intercept) | -0.823 | 0.404 | -2.038 | 0.042 | -1.614 | -0.032 |
| Multi-Month | activityModerately Active | -0.426 | 0.518 | -0.822 | 0.411 | -1.442 | 0.590 |
| Multi-Month | activityLess Active | -13.320 | 296.320 | -0.045 | 0.964 | -594.098 | 567.457 |
| Multi-Month | sexFemale | 1.076 | 0.558 | 1.926 | 0.054 | -0.019 | 2.170 |
| Multi-Month | sleepShorter Sleep | -0.231 | 0.482 | -0.479 | 0.632 | -1.175 | 0.713 |

#### Appendix 8: Cycle Synchrony

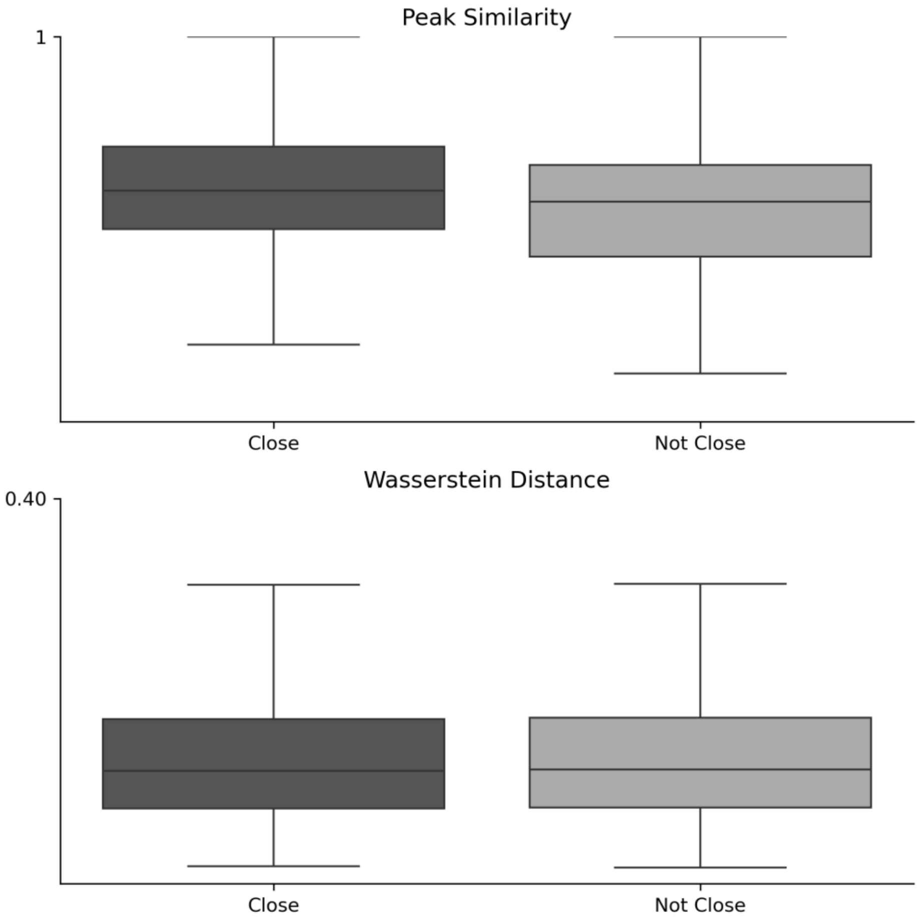


**Supplementary Figure 8: Spectral coherence between pairs of individuals.** Figure displays two metrics peak similarity and Wasserstein distance used for assessing the coherence between pairs of individuals with 'close' (i.e., friends, family, significant others) and 'not close' (i.e., strangers) relationships.

Individuals in the cohort were required to periodically complete social Network Surveys through which they listed 20 other individuals with whom they interacted with over the previous few months. This list often contained other participants in the study. To investigate physiological synchrony in heart rate cycles, cycles were compared between those with close relationships and those without close relationships in the cohort. The overall degree of "closeness" in a relationship was determined by how the individual defined others in their network and the frequency of contact between the two individuals. Family members (including two pairs of siblings identified), romantic partners, roommates, and individuals in contact on daily and weekly bases were defined as "Close"; on the other hand, those who did not fit into these categories (i.e., strangers, acquaintances, and friends with monthly or less than monthly frequencies of contact) were considered "Not Close".

Two nonparametric techniques Peak Similarity (PS) and Wasserstein Distance (WD) were employed to quantify the similarity between two individuals' periodogram peaks and overall shape. WD is a distance measurement between probability distributions, or in this case, two periodograms: a smaller WD indicates greater similarity between the periodograms. PS was determined by comparing the proportions of similar peaks within a pair. The individual with the greater number of peaks was set as a base for comparison, and the proportion of matching peaks to the other individual was calculated and regarded as the value of PS for that pair. If both individuals had the same number of peaks, the proportions and overall PS value were identical.

The results of this analysis are displayed in Supplementary Figure 7, visually depicting how those with closer relationships tended to have higher PS and very slightly higher WD values. Performing a Mann-Whitney U nonparametric statistical test on the PS and WD across the cohort, there was a statistically significant difference ($p = 0.004$) of PS values between the 'Close' and 'Not Close' groups. There was not a significant difference found for the WD ($p = 0.451$). These results indicate that individuals in closer relationships tended to have similar underlying heart rate cycles and are thus more environmentally similar.
